# A presynaptic spectrin network controls active zone assembly and neurotransmitter release

**DOI:** 10.1101/812032

**Authors:** Qi Wang, Lindsey Friend, Rosario Vicidomini, Tae Hee Han, Peter Nguyen, Chun-Yuan Ting, Mihaela Serpe

## Abstract

We have previously reported that *Drosophila* Tenectin (Tnc) recruits αPS2/βPS integrin to ensure structural and functional integrity at larval NMJs (Wang et al., 2018). In muscles, Tnc/integrin engages the spectrin network to regulate the size and architecture of synaptic boutons. In neurons, Tnc/integrin controls neurotransmitter release. Here we show that presynaptic Tnc/integrin modulates the synaptic accumulation of key active zone components, including the Ca^2+^ channel Cac and the active zone scaffold Brp. Presynaptic α-Spectrin appears to be both required and sufficient for the recruitment of Cac and Brp. We visualized the endogenous α-Spectrin and found that Tnc controls spectrin recruitment at synaptic terminals. Thus, Tnc/integrin anchors the presynaptic spectrin network and ensures the proper assembly and function of the active zones. Since pre- and postsynaptic Tnc/integrin limit each other, we hypothesize that this pathway links dynamic changes within the synaptic cleft to changes in synaptic structure and function.

## INTRODUCTION

The synaptic extracellular matrix (ECM) contains various synaptic organizers and diffusible molecules that establish and modify synaptic connections during development. ECM receptors, including integrins, transduce these signals by sensing the changes in the synaptic environment and orchestrating the appropriate changes in synapse structure and function (Chavis and Westbrook, 2001; Huang et al., 2006). Disruption of integrins impair synapse development and maturation, neurotransmission, and long-term synaptic plasticity (reviewed in (Park and Goda, 2016)). Integrins together with their ECM ligands form complexes with cytoskeletal proteins and active zone (AZ) components (Carlson et al., 2010; Nishimune et al., 2004; Sunderland et al., 2000). However, it remains unclear how these interactions could influence neurotransmitter release.

Spectrin, a principal component of the membrane skeleton, is critical for synaptic transmission (Featherstone et al., 2001; Goodman, 1999). Spectrin subunits function as docking platforms for membrane channels, neurotransmitter receptors and adhesion molecules (reviewed in (Machnicka et al., 2014)). For example, spectrins associate with presynaptic Ca^2+^ channels in synaptosome lysates (Carlson et al., 2010; Khanna et al., 2007a; Khanna et al., 2007b). Also, a spectrin-containing presynaptic web enables AZ function in the rodent CNS (Phillips et al., 2001). Recent super-resolution fluorescence microscopy uncovered periodic actin/spectrin lattices along axons, dendrites and spines in a broad range of neuronal cell types (He et al., 2016; Qu et al., 2017; Xu et al., 2013); however, at presynaptic and postsynaptic sites, these patterns were absent (Sidenstein et al., 2016).

We have previously found that *Drosophila* Tenectin (Tnc), an integrin ligand that accumulates in the synaptic cleft at the larval NMJ, modulates synapse development and function (Wang et al., 2018). Tnc recruits αPS2/βPS integrin and forms *cis* active complexes with distinct pre- and post-synaptic functions. In muscle, Tnc/integrin complexes regulate the size and architecture of synaptic boutons, partly through recruitment of the spectrin-based membrane skeleton. In motor neurons, Tnc/integrin complexes modulate neurotransmitter release. Since spectrin mutations showed similar neurotransmission deficits (Featherstone et al., 2001), we proposed that presynaptic Tnc/integrin functions by recruiting presynaptic spectrin (Wang et al., 2018).

Here we tested this hypothesis by investigating the presynaptic functions of Tnc/integrin complexes. We found that Tnc/integrin modulates the synaptic accumulation of key players in neurotransmitter release, including the voltage-gated Ca^2+^ channel, Cac (Macleod et al., 2006; Peng and Wu, 2007; Smith et al., 1996), and the AZ scaffold, Brp (Kittel et al., 2006; Wagh et al., 2006). We demonstrate that presynaptic α-Spectrin is both required and sufficient for Cac and Brp accumulation at the AZ. Using gene-editing and GFP-reconstitution approaches, we selectively labeled α-Spectrin in distinct synaptic compartments and revealed a Tnc-dependent presynaptic network. Together with our previous finding that postsynaptic Tnc/integrin recruits spectrin to modulate the size and architecture of synaptic boutons, this study identifies two spectrin-dependent integrin signaling pathways that coordinate synapse development and function.

## RESULTS

### Neuronal Tnc/integrin complexes regulate the recruitment of Ca^2+^ channels at the AZ

To facilitate our investigations of the Tnc role on neurotransmitter release, we first used the CRISPR/Cas9 technology to generate a *tnc*^*null*^ mutant, referred to here as *tnc*^*ko*^ (see Materials and Methods). This deletion removes ∼20kb of coding sequence, including both known *tnc* transcripts (Figure 1-figure supplement 1A and see details in Materials and Methods) (Syed et al., 2012). Similar to other *tnc* alleles, most *tnc*^*ko*^ animals died during pupa stages; the few adult escapers were sickly and had severe locomotion defects. Also, integrin recruitment was impaired at *tnc* mutant NMJs, but was partly restored by neuronal Tnc (Figure 1-figure supplement 1B-H) (Wang et al., 2018). The mEJP frequency was significantly reduced at *tnc*^*ko/Df*^ larval NMJs, a defect characteristic for *tnc* loss of function. In addition, *tnc*^*ko/Df*^ mutant NMJs had significantly reduced EJP amplitudes (from 35.42 ± 2.62 mV to 17.38 ± 2.05 mV, *p*<0.0001, Supplemental Table). The consistency and amplitude of *tnc*^*ko*^ phenotypes made this allele ideal for examining the presynaptic functions of Tnc.

**Figure 1.**
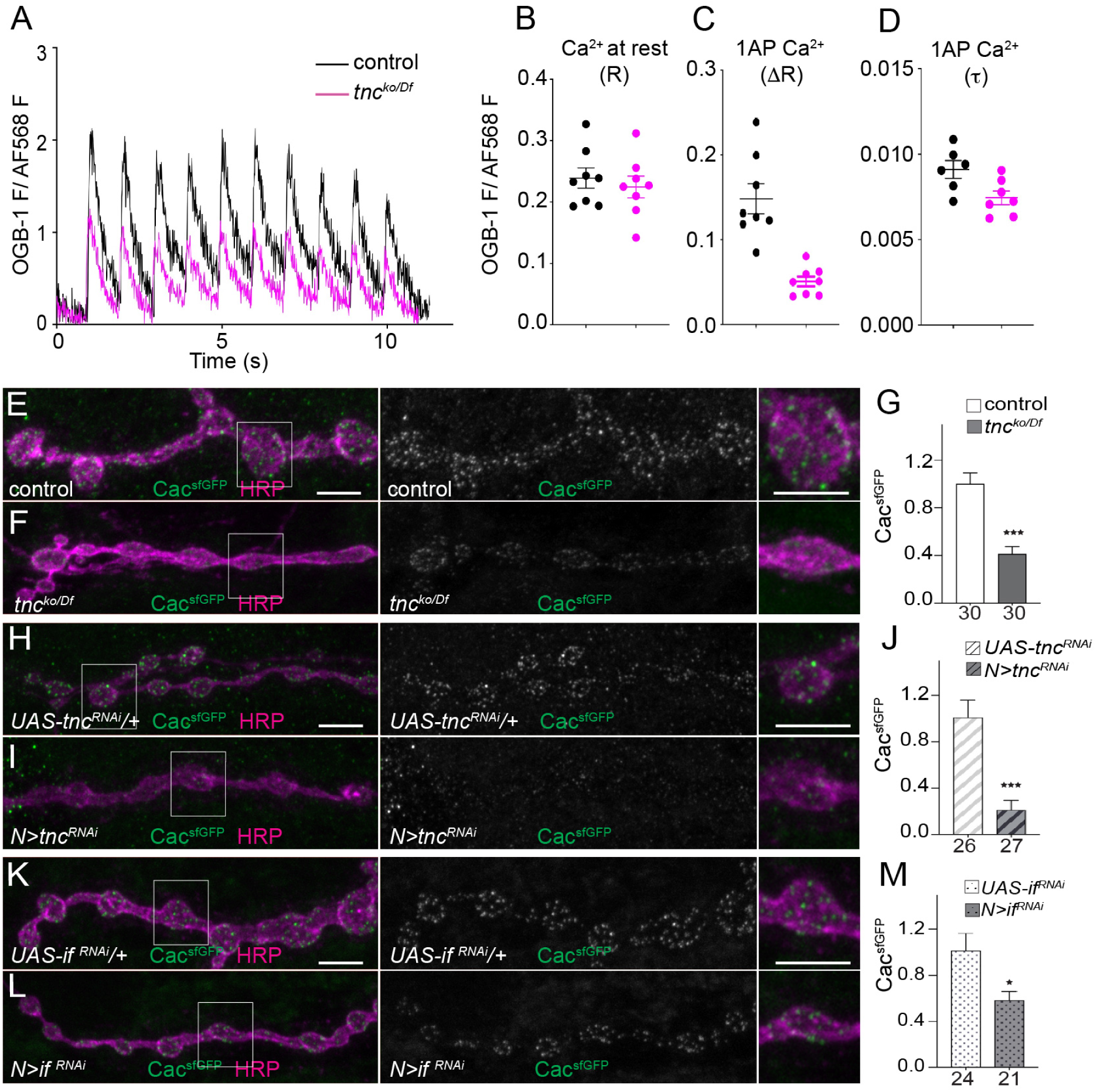
Tnc/integrin modulates the accumulation of Ca_V_2 Ca^2+^ channels at the AZ. (A) Single trial traces of changes in Ca^2+^-sensitive Oregon Green BAPTA-1 (OGB-1) fluorescence relative to Ca^2+^-insensitive Alexa Fluor 568 (AF568) fluorescence in response to stimuli (10 at 1Hz) applied to the hemisegment nerve. OGB-1 images collected at 100 frames per second. (B) Scatter plot of OGB-1/AF568 fluorescence prior to nerve stimulation, representing free Ca^2+^ levels in the cytosol ([Ca^2+^]_c_) at rest. Each circle represents a ratio (R) measurement from a synaptic terminal in a different larva. (C) Scatter plot of the amplitude (change in ratio: ΔR) of Ca^2+^ transients evoked by stimuli delivered at 1Hz. (D) Scatter plot of the decay time course (τ; reported in seconds) of Ca^2+^ transients evoked by 1Hz stimuli. (E-M) Confocal images of NMJs labeled for Cac^sfGFP^ (green) and HRP (magenta). Cac^sfGFP^ synaptic levels were strongly reduced in *tnc* mutants (*tnc*^*ko/Df*^), or in *tnc* or α*PS2* (*if*) integrin neuronal knockdown. Scale bars: 5 μm in (E, H and K). The number of NMJs examined is indicated below each bar. Bars indicated mean ±SEM. \**p*<0.05, \*\*\**p*<0.001. Genotypes: control (*Cac*^*sfGFP*^*/+); tnc*^*ko/Df*^ (*Cac*^*sfGFP*^*/+; tnc*^*ko*^*/Df(3R)BSC655*); *UAS-tnc*^*RNAi*^*/+* (*Cac*^*sfGFP*^*/+; UAS-tnc*^*RNAi*^*/+*); *N*>*tnc*^*RNAi*^ (*Cac*^*sfGFP*^*/BG380-Gal4; UAS-Dcr-2/UAS-tnc*^*RNAi*^); *UAS-if* ^*RNAi*^ */+* (*Cac*^*sfGFP*^*/+; UAS-if* ^*RNAi*^*/+*); *N*>*if* ^*RNAi*^ (*Cac*^*sfGFP*^*/BG380-Gal4; UAS-if RNAi/+*).

*tnc* mutants have elevated facilitation and significantly increased paired pulse ratio (PPR) (Wang et al., 2018), indicating that neurotransmitter release benefits from the accumulation of intracellular Ca^2+^, suggesting a presynaptic Ca^2+^ influx deficit. We tested this possibility using a Ca^2+^-sensitive fluorescent dye loaded into motor nerve terminals (Macleod, 2012). This dye fluoresces in proportion to free Ca^2+^ levels in the cytosol ([Ca^2+^]_c_); when loaded in constant proportion to a Ca^2+^ insensitive dye, it allows ratiometric comparisons between *tnc*^*ko/Df*^ and control terminals (ΔR) (Figure 1A-C). [Ca^2+^]_c_ at rest, estimated prior to stimulation, was no different in *tnc*^*ko/Df*^ relative to the control (Figure 1B). The amplitude of single AP evoked changes in [Ca^2+^]_c_ in response to 1Hz nerve stimulation was decreased from 0.148 ± 0.017 to 0.050 ± 0.005 in *tnc* mutants (n=8 for each group, *p*<0.001, Figure 1C). The single action potential decay time (τ) was only slightly decreased in *tnc* mutants (Figure 1D). Similar results were obtained with GCamP6s (not shown). These data reveal a significant deficit in Ca^2+^ entry in *tnc* terminals, but not in Ca^2+^ clearance. This deficit is consistent with the increased PPR observed at *tnc* mutant terminals (Wang et al., 2018).

Could Tnc play a role in the recruitment of Ca^2+^ channels at the AZ? Cac is the pore-forming subunit of the only Ca_V_2 Ca^2+^ channel required for neurotransmitter release at *Drosophila* NMJ (Macleod et al., 2006; Peng and Wu, 2007; Smith et al., 1996). Using an endogenously tagged *cac* allele, *cac*^*sfGFP*^ (Gratz et al., 2019), we found that Ca^2+^ channel levels were dramatically reduced at *tnc* NMJs (to 41.3 ± 6.1% compared to control, *p<*0.0001) (Figure 1E-G). Knockdown of neuronal Tnc similarly reduced the synaptic Cac^sfGFP^ levels to 21.5 ± 8.1% compared to the *UAS-tnc*^*RNAi*^ control (Figure 1H-J). To test whether Cac recruitment/stabilization at the AZs requires Tnc/Integrin complexes, we knocked down various integrin subunits in the *cac*^*sfGFP*^ background. Neuronal knockdown of *if*, the αPS2 integrin subunit, phenocopied the *tnc* mutants, and reduced the Cac^sfGFP^ synaptic signals to 58.5% compared to control (*p=*0.0165) (Figure 1K-M). Neuronal reduction of *mys*, the βPS integrin subunit, triggered a modest Cac^sfGFP^ reduction (not shown), probably because of redundancy with the neuronally abundant β*v* integrin subunit (Rohrbough et al., 2000). Together these results reveal an important role for the Tnc/integrin neuronal complexes in the clustering of Ca^2+^ channels at the AZ.

### Tnc/integrin complexes promote the synaptic accumulation of Brp

Reduced clusters of voltage-gated Ca^2+^ channels and depressed evoked vesicle release have also been reported for mutations in *brp*, the fly orthologue of ELKS/CAST scaffolds (Kittel et al., 2006). Indeed, synaptic Brp levels were reduced at *tnc*^*ko/Df*^ NMJs (to 58.3 ± 6.3% compared to control, n=24, *p*<0.0001) (Figure 2A-B, quantified in 2I). As described above, knockdown of Tnc in neurons, but not in muscle, triggered reduced Brp accumulation (to 75.4 ± 3% compared to control, n=25, *p*=0.0005) (Figure 2C-D and I). Also, neuronal knockdown of βPS significantly reduced the Brp intensity at synaptic terminals (to ∼80% compared to control, n=34, *p*=0.0013) (Figure 2E-F and I). Knockdown of αPS2 showed relatively normal Brp signal intensity in this experimental setting (Figure 2G-H and I), however, it did affect the Brp-marked structures (see below). This indicates that neuronal Tnc/integrin complexes influence the recruitment of Brp at the AZ.

**Figure 2.**
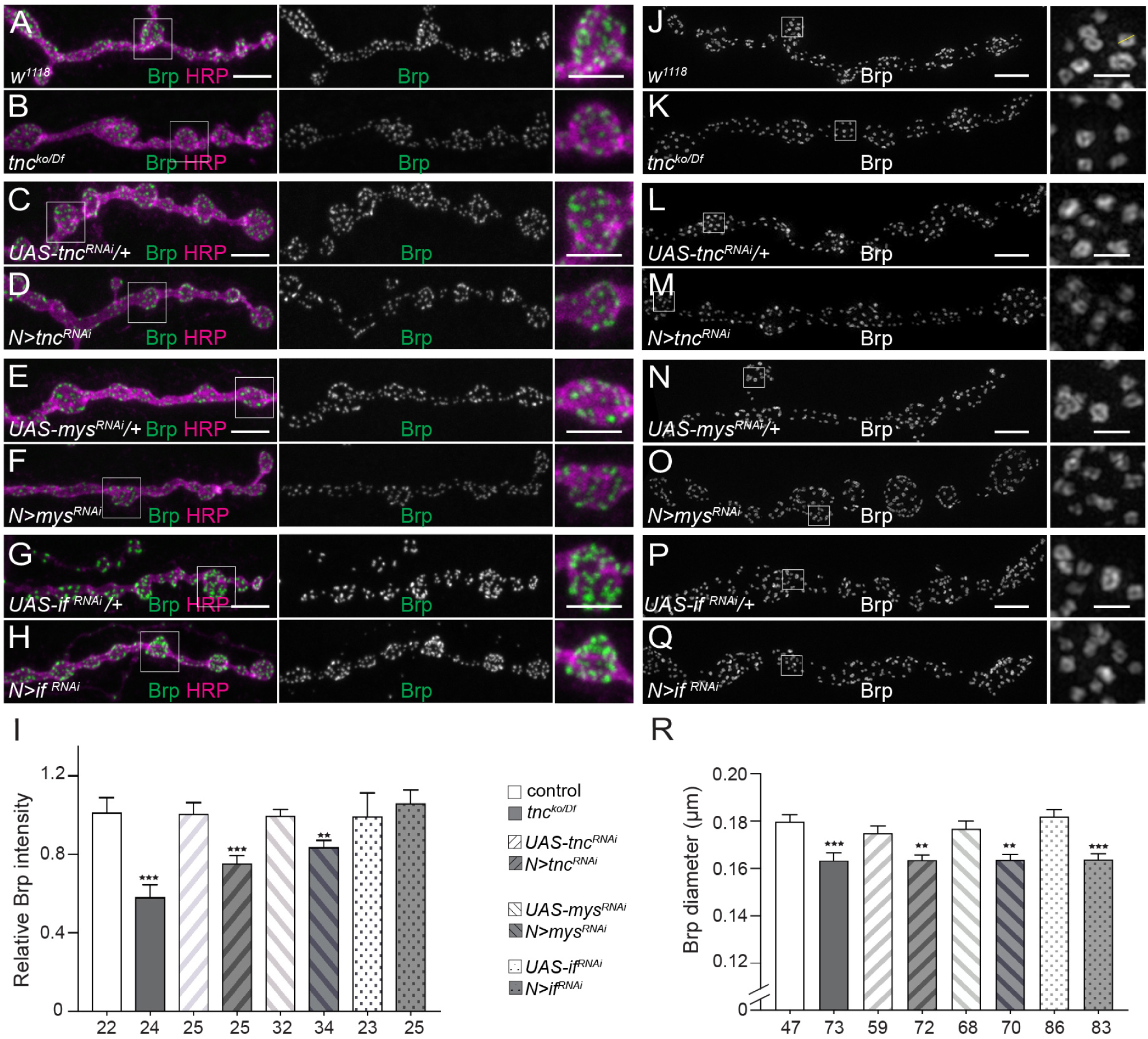
Neuronal Tnc/integrin regulates the AZ scaffold, Brp. (A-H) Confocal images of Brp-marked AZs in the indicated genotypes (quantified in I). *tnc*^*ko/Df*^ mutants or neuronal knockdown of *tnc* or *βPS* integrin diminished the synaptic Brp levels. (J-Q) 3D-SIM imaging of Brp-positive T-bar structures. Knockdown of *tnc* or α*PS2/βPS* subunits significantly reduced the T-bar diameters. Scale bars: 5 μm (A, C, E and G); 5 μm in (J, L, N and P); 200 nm in details. The number of examined NMJs (I) and individual synapses (R) is indicated below each bar. Bars indicated mean ±SEM. \*\**p*<0.01, \*\*\**p*<0.001. Genotypes: *tnc*^*ko/Df*^ (*tnc*^*ko*^*Df(3R)BSC655*); *N*>*tnc*^*RNAi*^ (*BG380/+; UAS-Dcr-2/UAS-tnc*^*RNAi*^); *N*> *mys*^*RNAi*^ (*BG380/+; UAS-Dcr-2/+; UAS-mys*^*RNAi*^*/+*); *N*> *if* ^*RNAi*^ ((*BG380/+; UAS-Dcr-2/UAS-if* ^*RNAi*^*/+*)

The anti-Brp monoclonal antibody Nc82 utilized here recognizes an epitope on the outer diameter of the T-bars and produces a ring-shaped signal when examined by super-resolution microscopy (Fouquet et al., 2009; Sulkowski et al., 2016). Using 3D structured illumination microscopy (3D-SIM), we measured the maximum diameter of the Brp-marked rings using a line plot across a single z slice (yellow line Figure 2J). The Brp diameter was slightly but significantly decreased at *tnc* NMJs (to 163.4 ± 3.3 nm, n=73, compared to control 178.8 ± 3.0 nm, n=47, *p*=0.0007) (Figure 2J-K and R). Neuronal Tnc-αPS2/βPS complexes were primarily responsible for this function, since Brp diameters in various neuronal knockdowns were similar to those observed at *tnc* mutant NMJs (*N*>*tnc*^*RNAi*^ 163.5 ± 2.2 nm, n=72; *N*>*mys*^*RNAi*^ 163.6 ± 2.4 nm, n=70; *N*>*if*^*RNAi*^ 166.9 ± 2.3 nm, n=83) (Figure 2 L-R). Together these data indicate that Tnc/integrin complexes influence the synaptic accumulation of Brp.

### The presynaptic spectrin network is critical for neurotransmitter release

Previous biochemical studies showed that presynaptic Ca^2+^ channels and AZ components form complexes with laminin/presynaptic integrins and cytoskeletal proteins, including α2/β2 spectrin in Torpedo electric organ synapses, which resemble NMJs (Carlson et al., 2010). In flies, *spectrin* mutations or neuronal knockdown of *α-Spec* and *β-Spec* disrupt presynaptic release similar to *tnc* and *integrin* knockdown (Featherstone et al., 2001; Pielage et al., 2005; Wang et al., 2018). Also, postsynaptic Tnc/Integrin recruits α-Spec and ensure bouton integrity (Wang et al., 2018), raising the possibility that presynaptic Tnc/integrin may also function by recruiting spectrin. We found that neuronal knockdown of α-Spec triggered a marked decrease of both synaptic Brp and Cac, to 63% (n=28, *p*=0.002) and 75% (*p*=0.014), respectively, compared with control (*UAS-α-Spec*^*RNAi*^*/+*, n=24) (Figure 3A-B). Also, *α-Spec* interacted genetically with *tnc*: trans-heterozygotes animals (*α-Spec/+, tnc/+*) exhibited mEJP and EJP deficits that resembled *tnc* loss of function, whereas individual heterozygote larvae (*α-Spec/+* or *tnc/+*) showed relatively normal recordings (Supplemental Table).

**Figure 3.**
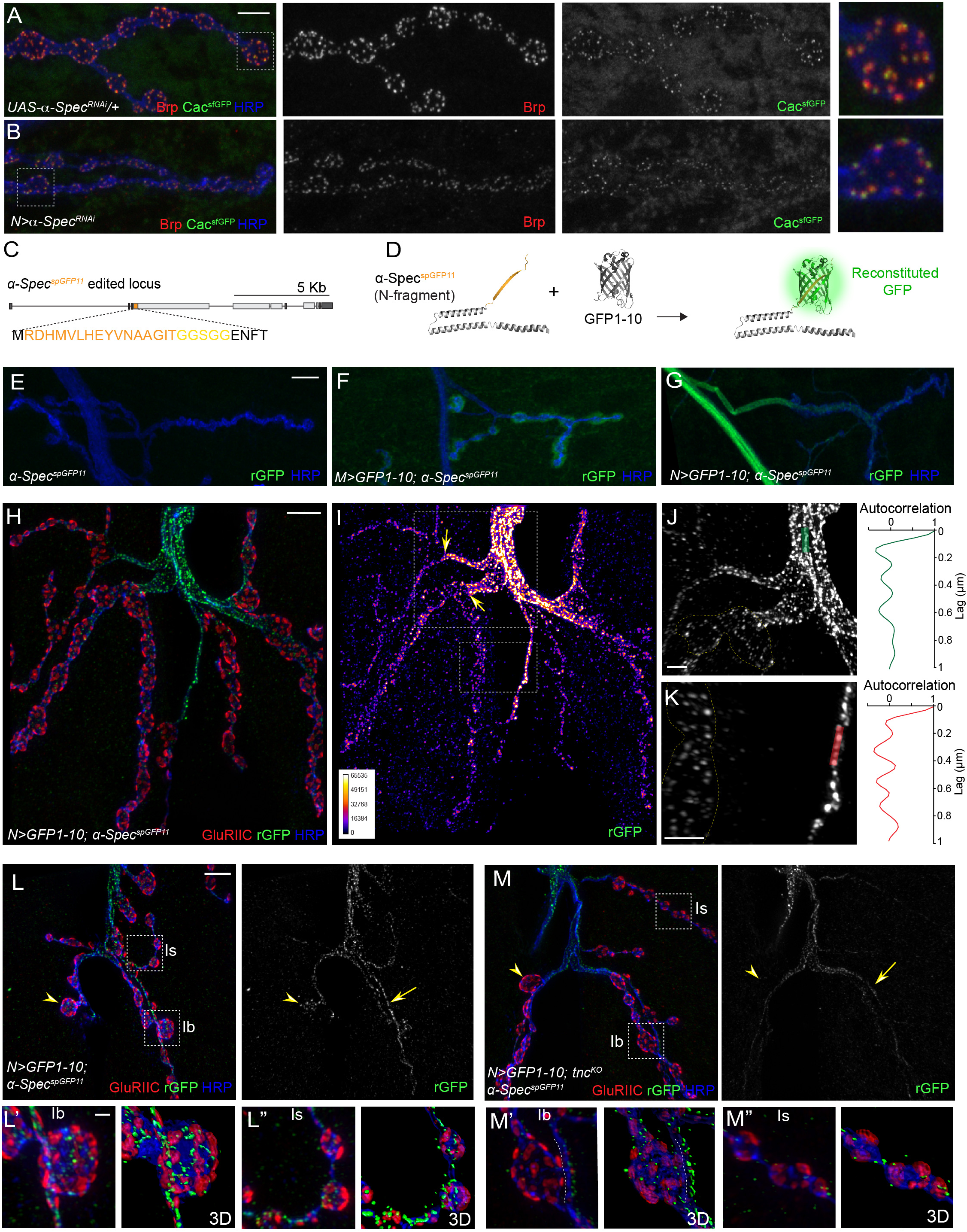
Neuronal Tnc stabilizes α-Spectrin at synaptic terminals. (A-B) Confocal images of NMJs labeled for Cac^sfGFP^ (green), Brp (red), and HRP (blue). Neuronal knockdown of α-Spectrin diminished the levels of both Brp and Cac^sfGFP^. (C-D) Diagram of spGFP11-tagged α-Spectrin and reconstitution of cytoplasmic GFP. (E-G) Confocal images of muscle 4 NMJs of third instar larvae of indicated genotypes labeled for reconstituted GFP (green) and HRP (blue). Expression of cytoplasmic GFP1-10 in *α-Spec*^*spGFP11*^ animals triggered compartment specific GFP reconstitution. (H-I) 3D-SIM images of muscle 12/13 NMJ labeled for GluRIIC (red), HRP (blue) and neuronally reconstituted GFP (green or Fire-lut). On the intensity scale (I), white represents peak intensity. The rGFP signals were intense along the axons but dropped at the NMJ (arrows). (J-K) Autocorrelation analysis along bundled (green line -J) or single (red line -K) axons captured the ∼190 nm periodicity of the spectrin lattice. (L-M) 3D-SIM images of muscle 4 NMJs and boutons labeled for neuronal rGFP (green), GluRIIC (red), and HRP (blue). The rGFP signals form interbouton bundles (arrows) and fan out in control boutons (arrowheads); these signals were reduced in the absence of Tnc. 3D reconstitution of Ib and Is terminals show a severe loss of spectrin in *tnc* mutant boutons. Scale bars: 10 μm (A and E), 5 μm (H and L), 2 μm (J and K) and 1 μm (L’). Genotypes: control (*α-Spec*^*spGFP11*^); *M*>*GFP1-10;α-Spec*^*spGFP11*^ *(G14-Gal4/UAS-GFP1-10; α-Spec*^*spGFP11*^*); N*>*GFP1-10;α-Spec*^*spGFP11*^ *(BG380-Gal4/+;UAS-GFP1-10/+; α-Spec*^*spGFP11*^); *N*>*GFP1-10;α-Spec*^*spGFP11*^,*tnc*^*ko*^ *(BG380-Gal4/+;UAS-GFP1-10/+; α-Spec*^*spGFP11*^,*tnc*^*ko*^).

To detect neuronal α-Spec, we employed a GFP reconstitution approach (Feinberg et al., 2008), whereby (a) one fragment of the split GFP, the 16 residue long spGFP11, was introduced into the *α-Spec* locus via CRISPR/Cas9, and (b) the second part, the spGFP1-10 fragment, was expressed intracellularly using the *Gal4/UAS* system (Brand and Perrimon, 1993) (Figure 3C-D, Figure 3-Supplement Figure 1, and Materials and Methods). Importantly, the *α-Spec*^*spGFP11*^ homozygous animals were viable and fertile, and had no observable defects, indicating that this insertion did not interfere with α-Spec function. Overexpression of spGFP1-10 in the muscle induced robust reconstituted GFP (rGFP) signals in *α-Spec*^*spGFP11*^ animals, but not in control (Figure 3E-F). Likewise, neuronal expression of spGFP1-10 in *α-Spec*^*spGFP11*^ animals produced a bright rGFP-positive lattice along the axons and progressively weaker rGFP signals at synaptic terminals (Figure 3G).

Recent super-resolution studies using overexpressed transgenes have uncovered a spectrin-based periodic membrane lattice in mammalian and *Drosophila* neurons (Qu et al., 2017; Xu et al., 2013). Using 3D-SIM, we also captured the periodicity of rGFP-marked endogenous α-Spec (Figure 3H-K). The rGFP signals were very strong along the motor neuron axons but dropped significantly where motor neuron arbors sink into the muscle cortex (arrows in Figure 3I). Autocorrelation analyses indicated that the lattice spacing was ∼200 nm (Figure 3J-K), consistent with previous measurements (He et al., 2016). Within the synaptic terminal, the rGFP-marked α-Spec fanned out and filled the synaptic boutons but coalesced into a bundle in the interbouton regions (Figure 3L). Interestingly, in the absence of Tnc, the rGFP-marked α-Spec signals remained bright along the axons but were drastically diminished at the synaptic terminals (Figure 3M, and Figure 3, Supplement Figure 1). These localized defects are consistent with the Tnc distribution, surrounding all the synaptic boutons (Wang et al., 2018). 3D reconstitution of type Ib and Is boutons showed robust rGFP staining in both terminals in control, but sparse bouton staining and reduced interbouton signals in *tnc* mutants. Quantification of the rGFP signals at synaptic terminals indicated a relative reduction by 40% (n=29, *p*<0.0001) at *tnc* mutant NMJs (Figure 3, Supplement Figure 1). Furthermore, neuronal knockdown of Tnc diminished the rGFP-marked α-Spec by 26% (n=17, *p=*0.034). Together our data indicate that presynaptic Tnc modulates the recruitment/stabilization of the spectrin lattice at synaptic terminals without affecting the lattice along the motor neuron axons.

### Presynaptic α-Spectrin functions downstream of Tnc to modulate AZ organization

If presynaptic, Tnc-dependent α-Spec modulates Cac and Brp recruitment, then excess of neuronal α-Spec should alleviate some of the *tnc* mutant deficits. Indeed, neuronal overexpression of α-Spec partly restored the Brp and Cac^sfGFP^ levels at *tnc* synapses (Figure 4A-C, quantified in D); these boutons remained small, a characteristic of *tnc* loss of function, but the levels of Brp and Cac were significantly increased compared with control. Thus, α-Spec is both required and sufficient for the recruitment of Cac and Brp at the AZs. Excess neuronal α-Spec caused substantial larval lethality (not shown), likely by disrupting axonal transport (Bennett and Lorenzo, 2013).

**Figure 4.**
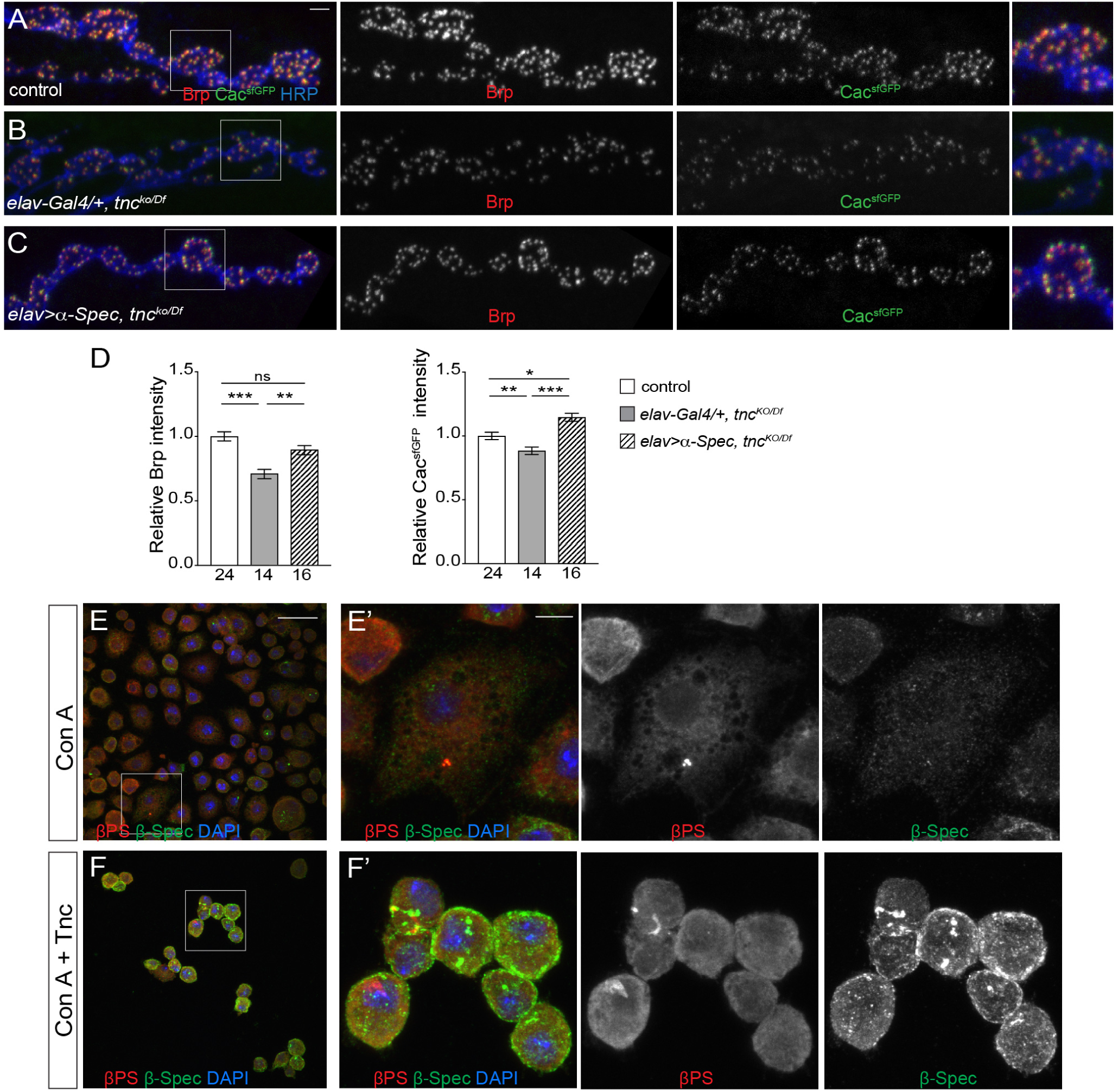
α-Spectrin regulates recruitment of Brp and Ca^2+^ channels at the AZ. (A-C) Confocal images of NMJs labeled for Cac^sfGFP^ (green), Brp (red), and HRP (blue) in third instar larvae of the indicated genotypes. Neuronal overexpression of α-Spec in *tnc* mutants partly restored the levels of Brp and Cac^sfGFP^ (quantified in D). (E-F) Confocal images of S2R+ cells spread on different substrates as indicated and stained for βPS (red), β-Spec (green) and DAPI (blue). Cells spread on Tnc coated surfaces show massive cortical accumulation of β-Spec (arrowheads). Scale bars: 10 μm (A-C), 30 μm (E-F) and 7 μm (E’-F’). The number of NMJs examined is indicated below each bar. Bars indicated mean ±SEM. ns -*p*>0.5, \**p*<0.05, \*\**p*<0.01, \*\*\**p*<0.001. Genotypes: control (*Cac*^*sfGFP*^*/Y); elav-Gal4*; *tnc*^*ko/Df*^ (*Cac*^*sfGFP*^*/Y; elav-Gal4, tnc*^*ko*^*/Df(3R)BSC655*); *elav*>*α-Spec*; *tnc*^*ko/Df*^ (*Cac*^*sfGFP*^*/Y; UAS-Myc-α-Spec/+; elav-Gal4, tnc*^*ko*^*/Df(3R)BSC655*).

To directly probe for Tnc-mediated spectrin recruitment at the cell membrane, we examined the behavior of cells spread on surfaces coated either with or without Tnc. As expected, S2R+ cells attached and spread outward on control coverslips coated with Concanavalin A (ConA) (Figure 4E). In contrast, when presented with the ConA/Tnc coated coverslips, the S2R+ cells rounded up and self-aggregated (Figure 4F). Notably, these round cells accumulated very high levels of β-Spec near the cell surface (Figure 4F, arrows), whereas control cells showed exclusively cytoplasmic distribution of β-Spec. The behavior of cells on ConA/Tnc coated surfaces is in stark contrast with that of cells presented with other RGD-containing ligands, such as fibronectin (Ribeiro et al., 2014); upon ligand binding, active integrins usually promote cell adhesion to substrates, formation of focal adhesion points, and ultimately enable cell migration. The exposure to Tnc triggered the opposite cell behavior, including the detachment of cells from surfaces, retraction of cell protrusions, and aggregation.

## DISCUSSION

The ECM and its receptors play critical roles in the assembly and function of the nervous system. In this study we reveal an unprecedented link between ECM/integrin complexes and spectrin-mediated AZ assembly and function. Our data are consistent with a model whereby neuronal Tnc/integrin complexes anchor the presynaptic spectrin network to ensure the proper clustering of Ca^2+^ channels and Brp scaffold at the AZs, thus enabling AZ functionality.

Several lines of evidence support this model. First, *tnc* mutants had electrophysiological deficits (*i.e.* paired pulse facilitation) consistent with reduced presynaptic Ca^2+^ entry (Figure 1 and (Wang et al., 2018)). Secondly, the Ca^2+^ channel levels were reduced by half at *tnc* terminals (Figure 1). Brp levels were similarly decreased, weakening the AZ structures and causing transmission deficits (Figure 2 and (Wang et al., 2018)). Third, neuronal knockdown of Tnc and αPS2/βPS integrin recapitulated these defects, indicating that presynaptic Tnc/integrin complexes function to anchor the Ca^2+^ channels and organize the AZ. Fourth, knockdown of neuronal α-Spec diminished synaptic Ca^2+^ channels and Brp levels, whereas overexpression of α-Spec partly restored their accumulation at *tnc* mutant terminals; thus, presynaptic α-Spec is both required and sufficient for clustering of Ca^2+^ channels and Brp at AZ. Fifth, the presynaptic α-Spec network was drastically diminished at *tnc* mutant boutons, but appeared independent of Tnc along the motor neuron axons (Figure 3). We have previously demonstrated a similar Tnc-dependent recruitment of postsynaptic spectrin at larval NMJ (Wang et al., 2018). In both cases, Tnc and αPS2/βPS integrin form *cis* active complexes which may stabilize the pre- and post-synaptic membranes. Sixth, in the presence of Tnc, S2 cells accumulated integrin and spectrin at their membranes, but didn’t migrate and didn’t form focal adhesion points; instead, these cells shrunk their membranes and rounded up (Figure 4).

Since peaks of ecdysone-controlled Tnc (Fraichard et al., 2006) coincide with formation of new boutons, Tnc may play a major role in the restructuring of the synaptic structures. Mucins appear to be particularly effective at bending cell membranes (Shurer et al., 2019). At synaptic terminals, this cell behavior may facilitate formation of round boutons, with minimal cell surface, thus concentrating the AZ components. In the muscle, Tnc/integrin recruits postsynaptic spectrin to control the structural integrity of synaptic boutons (Wang et al., 2018). Thus, the interplay between Tnc/integrin and spectrin may regulate the dynamic remodeling of synaptic terminals and balance growth with bouton formation/stabilization in the developing motor neuron arbors.

It has been long established that spectrins and integrins play important roles in the synaptic transmission (Chavis and Westbrook, 2001; Goodman, 1999), including the gating of neurotransmitter release (Featherstone et al., 2001; Huang et al., 2006). Presynaptic integrin and spectrin subtypes have been directly implicated in interactions with voltage-gated Ca^2+^ channels and AZ components (Carlson et al., 2010; Garcia-Caballero et al., 2018; Khanna et al., 2007a; Khanna et al., 2007b). Nonetheless, a role for ligand-activated integrin in the synaptic recruitment of spectrin has not been previously reported. Biochemical studies in various systems uncovered complexes containing ECM components (*i.e.* synaptic laminin) and non-erythroid spectrin together with voltage-gated Ca^2+^ channels and AZ components (Carlson et al., 2010; Sunderland et al., 2000). It was proposed that such complexes, containing presynaptic voltage-gated Ca^2+^ channels, synaptic laminin, α2/β2 spectrins, and two AZ components (Bassoon and Piccolo), may couple channel-anchoring to assembly and/or stabilization of AZs. Our results suggest that such coordination may lie within the synaptic cleft: Dynamic changes in the ECM could be transduced via ligand-activated integrin and spectrin to mobilize the Ca^2+^ channels and other AZ components and organize the AZs.

To our knowledge, this is the first visualization of the endogenous presynaptic spectrin mesh in a fully functional synapse. Previous studies used overexpressed GFP-tagged transgenes or RNAi depletion of the postsynaptic spectrin pool (He et al., 2016; Pielage et al., 2005). Such settings create significant disruptions of synapse development and function, affecting spectrin distribution. Instead, our study captures the endogenous spectrin distribution, validating previous reports of a periodic spectrin lattice along the axons (D’Este et al., 2015; He et al., 2016; Zhong et al., 2014). A step reduction of the spectrin network marks the entry point of the motor neuron in the muscle cortex (Figure 3 and (Pielage et al., 2005)). An expansion of the lattice occurs in each synaptic bouton (Figure 3 and (Han et al., 2017; Sidenstein et al., 2016)). Importantly, Tnc controls the spectrin lattice locally, within the synaptic boutons, but not along the axon, indicating that Tnc-dependent functions are localized to the tetrapartite synapse. Also, additional means of anchoring spectrin along the inter-boutons must exist, for example Teneurins connect the spectrin cytoskeleton via trans-synaptic interactions (Mosca et al., 2012).

In the future it will be interesting to examine how Tnc/integrin complexes engage the spectrin-based lattice. Candidate intermediates include actin filaments or Dlg/MAGUK isoforms which could both bind integrin and cluster presynaptic voltage-gated Ca^2+^ channels (Astorga et al., 2016; Beumer et al., 2002; Guzman et al., 2019; Ruiz-Canada and Budnik, 2006). Integrin activation generally promotes formation of focal adhesion complexes and cell migration. Tnc triggers the opposite effect: Cells become round and compact and accumulate high levels of subcortical spectrin (Fig. 4). This is reminiscent of erythrocytes that maintain their shape and function through the membrane associated spectrin skeleton (Machnicka et al., 2014). Interestingly, several patients with acanthocytosis (spiky erythrocytes) caused by spectrin tetramerization deficits also present nervous system abnormalities, underscoring the relevance of spectrin network for normal neuronal development and function (Orlacchio et al., 2007). Together with our previous finding that postsynaptic Tnc/integrin recruits spectrin to modulate the size and architecture of synaptic boutons, this study identifies two spectrin-dependent integrin signaling pathways that mediate synapse development and function. While Tnc does not have a direct mammalian homolog, many synaptic cleft proteins have mucin domains and RGD motifs (*i.e.* Agrin (McMahan et al., 1992)), suggesting that similar ECM–integrin interactions may anchor spectrins to modulate synapse assembly and regulation.

## AKNOWLEDGMENTS

Q.W., L.F., R.V., T.H.H, P.N., and M.S. were supported by Intramural Program of the National Institutes of Health, *Eunice Kennedy Shriver* National Institute of Child Health and Human Development, grants ZIA HD008914 and ZIA HD008869 awarded to M.S. We thank Claire Thomas for generously sharing spectrin transgenes and antibodies and for comments on this manuscript. We thank Kate O’Connor-Giles for the *cac*^*sfGFP-N*^ flies. We thank Tom Brody for discussions on this manuscript and Greg MacLeod for help with Ca^2+^ measurements. We are grateful to Vincent Schram, Chad Williams and Daniela Malide for support with confocal and super-resolution imaging. We also thank the Bloomington Stock Center at Indiana University for fly stocks and the Developmental Studies Hybridoma Bank at the University of Iowa for antibodies.

## MATERIALS AND METHODS

### Fly stocks

The *tnc*^*KO*^ and *α-Spec*^*spGFP11*^ alleles were generated using classic CRISPR/Cas9 methodology as previously described (Gratz et al., 2015). Briefly, a pair of gRNAs were injected in *y sc v; [nos-Cas9]attP40/CyO* stock (Ren et al., 2013) alone (for *tnc*) or together with a donor (in the case of *α-Spec*), followed by germline transformation (Rainbow transgenics).

A series of unmarked *tnc* deletions were isolated and molecularly characterized by PCR from genomic DNA (QuickExtractDNA, Epicentre) and sequencing. Putative genetic null alleles have been isolated and confirmed by sequence analysis; three individual *tnc* deletions were identified. The primers used for gRNAs, PCR and sequencing were as follows:

tnc-1-sense: CTTCGAGACACGCCGGTTTGAGGG

tnc-1-antisense: AAACCCCTCAAACCGGCGTGTCTC

tnc-2-sense: CTTCGCAATAGAAATCATAGAGCC

tnc-2-antisense: AAACGGCTCTATGATTTCTATTGC

Tnc-16636F: GTGGTGGTTTCGGTATTTGG

Tnc-37537R: CTGTGGTGTAGTGTGGTGTG

The last two Tnc-F/R primers are predicted to amplify a 21kb product from control animals and ∼700 bp from *tnc*^*null*^. Line *tnc*^*ko13-2*^ missing 20kb (25,001,139 −25,021,503), including the entire coding sequence, was selected as *tnc*^*null*^ and referred here as *tnc*^*ko*^.

For editing *α-Spec* and addition of the spGFP11 sequence, 5’ and 3’ homologous regions (HR) were PCR amplified and a donor plasmid was assembled from (a) 5’HR BglII-BspHI fragment, (b) a BspHI-BamHI synthesized fragment (that includes the spGFP11 insertion right after the start, as indicated in Figure 3C, and restriction sites for further addition of the scarless GMR-3xP3-dsRed cassette (Gratz et al., 2015)), and (c) 3’HR BamHI-HindIII fragment. The primers used for gRNAs, PCR and sequencing were as follows:

α-Spectrin-up sense: CTTCGTTGATATTGAAAAGCTGTAA

α-Spectrin-up antisense: AAACTTACAGCTTTTCAATATCAAC

α-Spectrin-down sense: CTTCGCGGCATCGCGCTTGAAGTAC

α-Spectrin-down antisense: AAACGTACTTCAAGCGCGATGCCGC

α-Spectrin 5’HR-F: 5’GCGAATGCATCTAGATAGATCTGGCCGAGCTGCTTCGAAAAG

α-Spectrin 5’HR-R: 5’ TGGAAAAAACAAAATCATGAAACTGAATGGCTGACTATCGTA

α-Spectrin 3’HR-F: 5’GAACCAGCGAAATGGAGAACTTTACACCCAAAGAG

α-Spectrin 3’HR-R: 5’CAAAGATGTCTAAAGCCTTGATCTTCTCCTCCTGG

Upon germline transformation, a series of GMR-3xP3-dsRed-marked insertions were isolated, balanced and molecularly characterized. The fluorescently visible marker was next removed by PiggyBac transposase (Gratz et al., 2015), and the editing events further confirmed by PCR and sequencing, using the following primers:

Spec-5HR-1F: AGGAGTTAGCCACCAAAGTGT

Spec-5HR-1R: TGGGCGCTGGATTGTTTTTAAT

Spec-3HR-2F: AGACAAGTAAACGAACCAGCGA

Spec-3HR-2R: AATTAGCGCCTCCACAGAGT

Spec-CDS-3F: TCATCGACCTGAGCAACAGT

Spec-CDS-3R: GAGGCGGACCTGAATCGAC

GMR-DsRed-F: GTCGAGGGTTCGAAATCGATAAG

GMR-DsRed-R: GGCCGCGACTCTAGATCATAA

Several independent *α-Spec*^*spGFP11*^ lines were isolated; all were homozygous viable and appeared healthy, indicating that this insertion does not interfere with the function of α-Spectrin.

The spGFP1-10 cytoplasmic cassette (Feinberg et al., 2008) was cloned in plasmid pJFRC7-20XUAS-IVS-mCD8::GFP, a gift from Gerald Rubin (Addgene #26220)(Pfeiffer et al., 2010), and introduced at docking site VK18 by Phi31 integrase-mediated germline transformation to generate *20xUAS-spGFP1-10* (BestGene Inc.) (Venken et al., 2006).

Other fly stocks were used in this study were: *UAS-tnc, UAS-Dcr-2,UAS-tnc*^*RNAi*^, *UAS-mys*^*RNAi*^, *UAS-if*^*RNAi*^, *Df(3R)BSC655* (Wang et al., 2018); *α-Spec*^*rg41*^ (BSC 31999); *UAS-Myc::α-Spec* (from Claire H. Thomas); *Cac*^*sfGFP-N*^ (from Kate M. O’Connor-Giles)(Gratz et al., 2019). The *BG380-Gal4, 24B-Gal4, elav-Gal4*, and *G14-Gal4* drivers were previously described.

### Immunohistochemistry

Wandering third instar larvae of various genotypes were dissected as previously described in ice-cooled Ca^2+^-free HL-3 solution (70 mM NaCl, 5 mM KCl, 20 mM MgCl_2_, 10 mM NaHCO_3_, 5 mM trehalose, 5 mM HEPES, 115 mM sucrose) (Budnik et al., 2006; Stewart et al., 1994). The samples were fixed in 4% paraformaldehyde (PFA) (Polysciences, Inc.) for 15 min, washed in PBS containing 0.5% Triton X-100, then incubated with specific antibodies. The following primary antibodies were used: anti-Brp (Nc82, 1:200, Developmental Studies Hybridoma Bank); anti-βPS integrin (CF.6G11, 1:10, Developmental Studies Hybridoma Bank); anti-GFP1-20 (GFP-20, 1:500, Sigma-Aldrich); anti-GFP (FluoTag-X4 coupled with Atto 488, 1:500, NanoTag Biotechnologies); Cy5-conjugated goat anti-HRP, (1:2000, Jackson ImmunoResearch Laboratories, Inc.). The Alexa Fluor 488-, Alexa Fluor 568-, and Alexa Fluor 647-conjugated secondary antibodies (Invitrogen) were used at 1:200. All samples were mounted in ProLong Gold (Invitrogen).

Samples of different genotypes were processed simultaneously and imaged under same confocal settings with a laser scanning confocal microscope (CarlZeiss LSM780). All images were collected as 0.2μm optical sections and the *z*-stacks were analyzed with Imaris software (Bitplane). To quantify fluorescence intensities, synaptic ROI areas surrounding anti-HRP immunoreactivities were selected and the signals measured individually at NMJs (muscle 4, segment A3) from ten or more different larvae for each genotype (number of samples is indicated in the graph bar). The signal intensities were calculated relative to HRP volume and subsequently normalized to control. Each experiment was performed at least three times. The numbers of total samples analyzed are indicated inside the bars. Statistical significances were performed by Prism8 using two-tailed unpaired t tests for two conditions or one-way ANOVA followed by Tukey test for three or more conditions. Error bars in all graphs indicate standard deviation ±SEM. \*\*\**p*<0.001, \*\**p*<0.005, \**p*<0.05, ns-*p*>0.05.

*Drosophila* S2R+ cells were used for expression and production of recombinant Tnc as previously described (Wang et al., 2018). For cell spreading assays, coverslips were precoated with Concanavalin A (Con A) (0.05mg/ml, Invitrogen) alone or with Con A+Tnc (Con A −0.05mg/ml plus Tnc −15μg/ml) at room temperature for 2 hours, washed and then incubated with S2R+ cells at room temperature for 1 hour. Excess cells were washed with fresh media, then attached cells were fixed in 4% PFA and stained for anti-β-Spectrin (rabbit polyclonal, 1:200, a gift from Claire H. Thomas) and anti-βPS integrin (CF.6G11, 1:10, Developmental Studies Hybridoma Bank). Alexa Fluor 488- and Alexa Fluor 568-conjugated secondary antibodies (Invitrogen) were used at 1:200. The samples were mounted in SlowFade Gold Antifade reagent with DAPI (Invitrogen), imaged by confocal microscopy and analyzed with Imaris software (Bitplane).

### Super resolution (3D-SIM) imaging

Super resolution imaging was used to study the structure of Brp rings and reconstituted *α-Spec*^*spGFP11*^ at the synaptic terminals. Images were collected by a Carl Zeiss Elyra PS1 inverted microscope with a Plan-Apo 100X (1.46 NA) oil immersion objective lens and an EM-CCD Andor iXon 885 camera. Raw images were acquired from various NMJs as specified at the A3 segment with ×5 phases at ×3 angles in the 1024 pixels resolution per plane, the interval for z-section stacks is 100 nm. All raw images were processed and reconstructed in 3D using Zen Black 2010 software (Carl Zeiss) with appropriate parameters.

For measuring the size of Brp rings, reconstructed images were further analyzed by image J. The fluorescent intensity of either side for an isolated Brp ring was measured by Plot Profile tool, and the distance between the two intensity peaks was calculated as diameter of Brp rings. At least 5 animals for each genotype were quantified, the exact number of quantified Brp rings was indicated in these graphs. Statistical analyses were performed with Prism8 by using the Student’s t-test.

To estimate the periodic skeletal structure of the reconstituted *α-Spec*^*spGFP11*^, processed 3D-SIM images were further analyzed by ImageJ. ROIs tracing the lattice along the axons were selected and the Plot Profile values (pixel, fluorescence intensity) were exported as Microsoft Excel file. Autocorrelation analysis of the listed values was performed using the following formula:

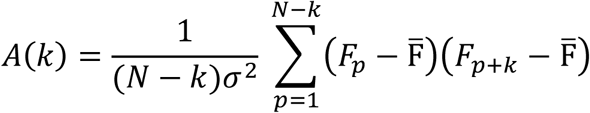

where *N* = total number of pixels of thePlotProfile, *p* = specific pixel, *F*_*p*_= Fluorescence value of the specific pixel *p*, 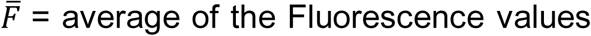, *σ*^2^= variance of the Fluorescence values, and *k*= lag = 1pixel = 0.025 nm.

### Electrophysiology

The standard larval body wall muscle preparation developed by Jan and Jan (1976) (Jan and Jan, 1976) was used for electrophysiological recordings. Wandering third instar larvae were dissected and washed in HL-3 using a custom microscope stage system. All recordings were performed in HL-3 saline (Stewart et al., 1994) containing 0.5 mM CaCl_2_. The nerve roots were cut near the exiting site of the ventral nerve cord so that the motor nerve could be picked up by a suction electrode. Intracellular recordings were made from muscle 6, abdominal segment 3 and 4. Data were used when the input resistance of the muscle was 5-15 mΩ and the resting membrane potential was between −60 mV and −70 mV. The input resistance of the recording microelectrode (backfilled with 3 M KCl) ranged from 20 to 25 mΩ. Muscle synaptic potentials were recorded using Axon Clamp 2B amplifier (Axon Instruments) and analyzed using pClamp 10 software. Spontaneous miniature excitatory junction potentials (mEJPs) were recorded in the absence of any stimulation. To calculate mEJP mean amplitudes, 50–100 events from each muscle were measured and averaged using the Mini Analysis program (Synaptosoft). Minis with a slow rise or falling time arising from neighboring electrically coupled muscles were excluded. Evoked EJPs were recorded following supra-threshold stimuli (200 μsec) to the appropriate segmental nerve with a suction electrode. Ten to fifteen EJPs evoked by low frequency of stimulation (0.1 Hz) were averaged. Corrected EJP amplitude was calculated using the following formula: E[Ln[E/(E -recorded EJP)]], where E is the difference between reversal potential and resting potential. The reversal potential used in this correction was 0 mV. ANOVA statistical analysis were preformed using Prism8 software. Data are presented as mean ±SEM.

### Presynaptic Ca^2+^ imaging

Cytosolic Ca^2+^ levels were monitored through the fluorescence of a Ca^2+^-sensitive dye (Oregon-Green BAPTA-1; OGB-1) relative to a Ca^2+^-insensitive dye (Alexa Fluor 568; AF568); both of which were loaded into motor neuron terminals using the forward-filling technique as previously described (Macleod, 2012). Segment nerves were forward-filled with 10,000 MW dextran-conjugated OGB-1, in constant ratio with 10,000 MW dextran-conjugated AF568. Fluorescence imaging was performed through a water-dipping 60X 1.1 NA Olympus objective fitted to an upright Olympus BX51W1 microscope. Fluorescence was excited using a CoolLED pE-4000 light source (OGB-1: 483/32 nm; AF568: 550/15 nm). Emitted light (OGB-1: 525/84 nm; AF568: 605/52 nm) was captured by an Andor Zyla sCMOS camera running at 100 frames-per-second (2×2 binning, 8 msec exposures). AF568 fluorescence images were captured immediately before and after the stimulus protocol to provide ratio information. While larvae were dissected and incubated in Schneider’s insect medium, this medium was replaced with HL6 at least 20 minutes prior to imaging. HL6 was supplemented with 0.5 mM Ca^2+^, 20 mM Mg^2+^, and 7 mM L-glutamic acid (Macleod et al., 2004). Segmental nerves were stimulated wherein each fluorescence transient is the result of an impulse of approximately 1.5 volts applied to the nerve for 0.4 ms. The background fluorescence was subtracted from each image and the average pixel intensity was measured within a region-of-interest containing 2-5 non-terminal boutons using NIS-Elements AR software (Nikon). Fluorescence intensity traces were further processed in ImageJ. OGB-1 fluorescence was imaged for 5 seconds prior to the first stimulus and was used to estimate the OGB-1 bleach trend, which was then numerically removed from the entire trace. Ca^2+^ levels are expressed as the fluorescence ratio of OGB-1 to AF568 (ΔR). Fluorescence transients corresponding to the action potentials evoked at 1 Hz were numerically averaged into a single trace and used to calculate peak amplitude and the decay time constant (τ). Differences between *tnc*^*KO/Df*^ and control (*w*^*1118*^) were analyzed using the Students T-test.

## KEY RESOURCE TABLE

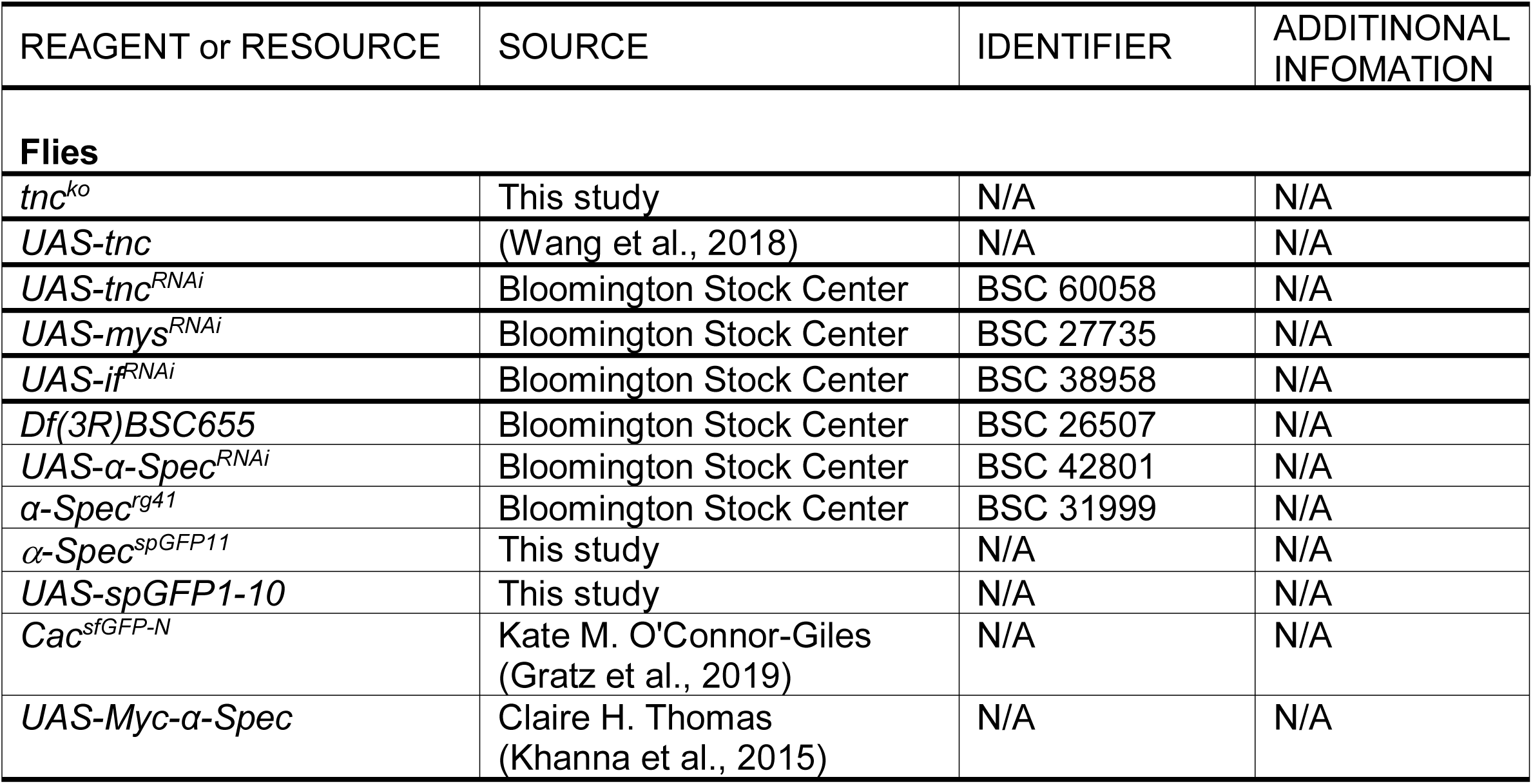
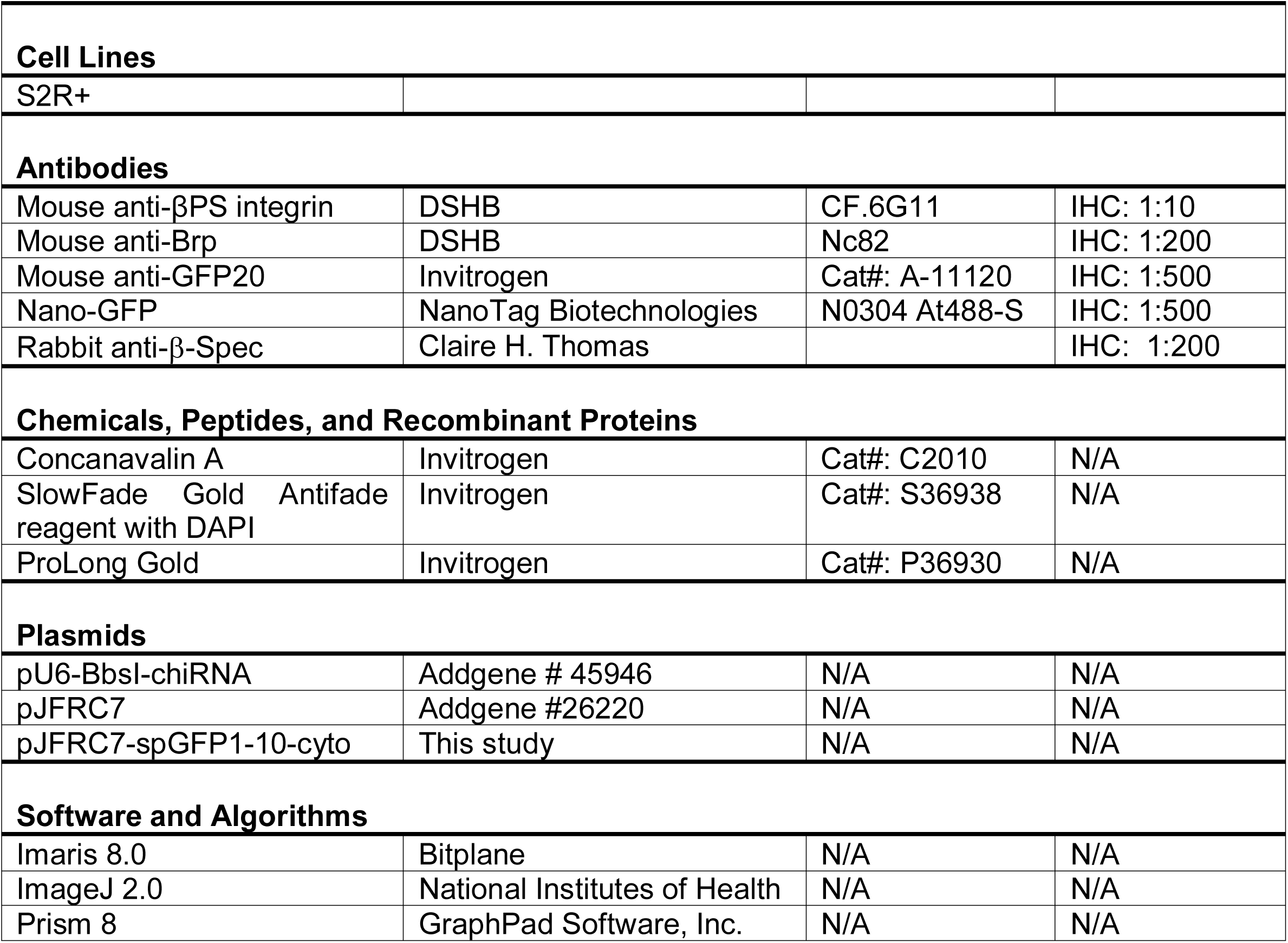

## Supplemental Table

| Relevant figure | Genotype | EJP Amp (mV) | mEJP Freq (Hz) | mEJP Amp (mV) | Vm<sub>rest</sub> | n |
| --- | --- | --- | --- | --- | --- | --- |
| Figure 1 and 3 | $w^{1118}$ | 35.425 ± 2.624 | 2.687 ± 0.308 | 1.198 ± 0.098 | 60.220 ± 0.364 | 10 |
| Figure 1 and 3 | $tnc^{ko/Df}$ | 17.378 ± 2.054 | 2.541 ± 0.173 | 1.033 ± 0.077 | 62.180 ± 1.506 | 11 |
| Figure 3 | $tnc^{ko/+}$ | 24.932 ± 1.035 | 2.414 ± 0.158 | 1.099 ± 0.139 | 63.220 ± 1.372 | 12 |
| Figure 3 | α-Spec/+ | 29.971 ± 1.277 | 2.802 ± 0.208 | 1.164 ± 0.091 | 61.700 ± 0.8172 | 10 |
| Figure 3 | $tnc^{ko/+}$; α-Spec/+ | 20.277 ± 1.278 | 2.138 ± 0.123 | 0.900 ± 0.106 | 63.090 ± 1.665 | 10 |

---

**Figure 1- figure supplement 1.**
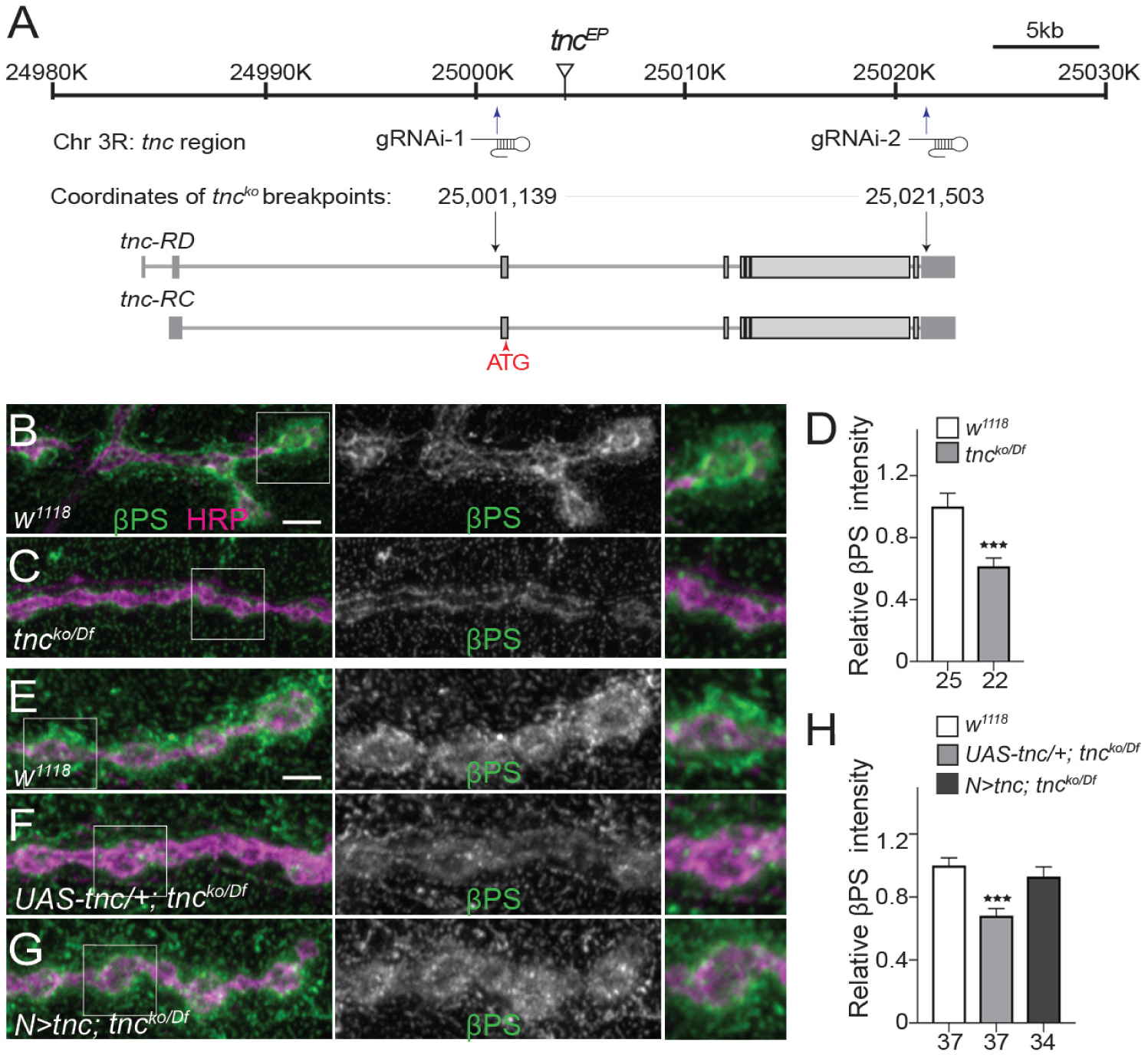
Generation and characterization of *tnc*^*null*^ mutant *(tnc*^*ko*^). (A) Diagram indicating the breakpoints of *tnc*^*ko*^ deletion. (B-G) Confocal images of third instar NMJs of indicated genotypes stained for βPS integrin (green) and HRP (magenta) (quantified in D-H). *tnc*^*ko/Df*^ mutants have reduced βPS synaptic levels that could be partly restored by neuronal expression of Tnc. Scale bars: 5 μm. The number of NMJs examined is indicated below each bar. Bars indicated mean ±SEM. \*\*\**p*<0.001. Genotypes: *tnc*^*ko/Df*^ (*tnc*^*ko*^*/Df(3R)BSC655*); *UAS-tnc/+; tnc*^*ko/Df*^ (*UAS-tnc/+*; *tnc*^*ko*^*/Df(3R)BSC655*); *N*>*tnc; tnc*^*ko/Df*^ (*UAS-tnc/+*; *tnc*^*ko*^*/Df(3R)BSC655, elav-Gal4*).

**Figure 3 - figure supplement 1.**
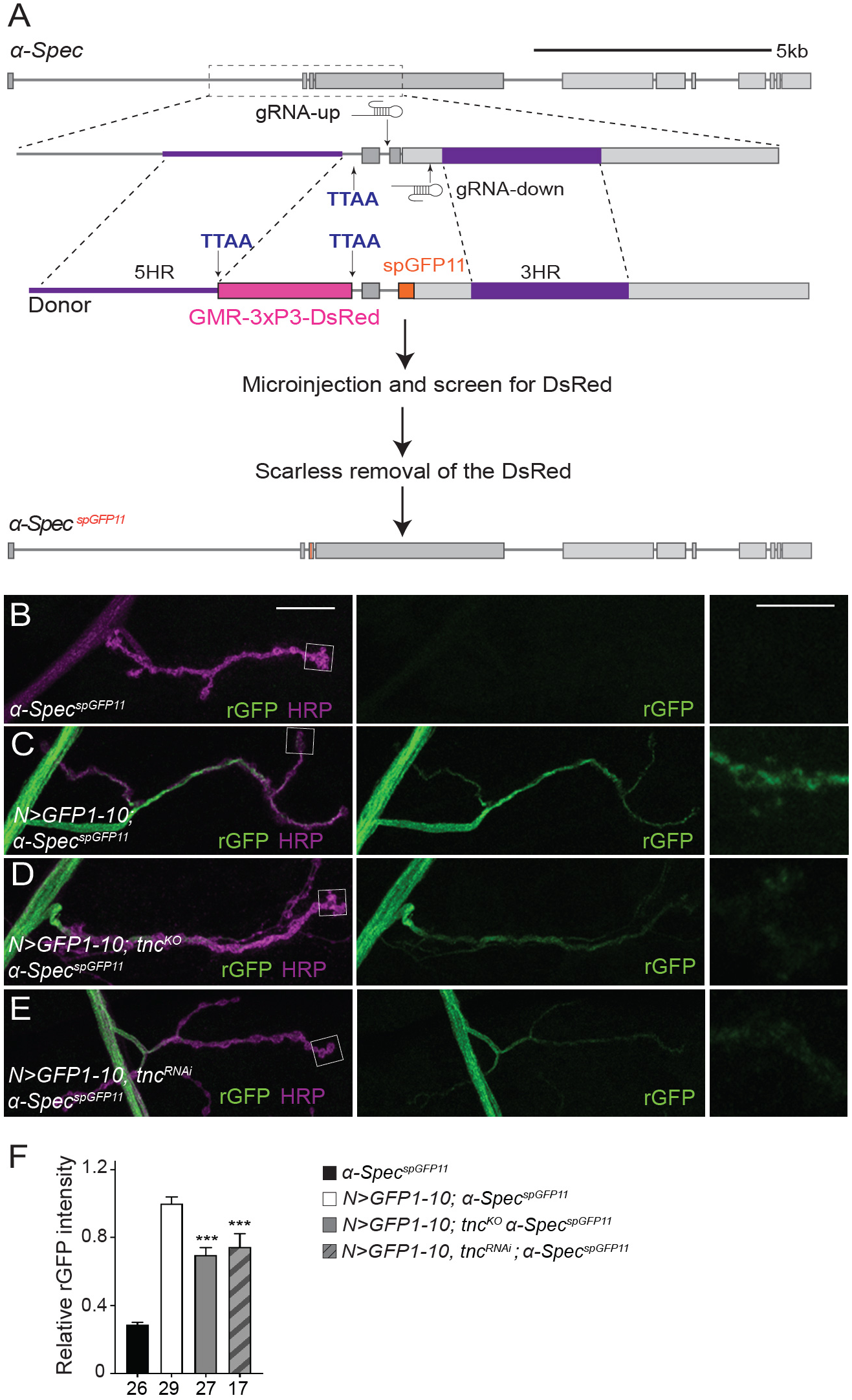
(A) Generation of scarless-edited *α-Spec*^*spGFP11*^. (B-E) Confocal images of NMJs labeled for neuron reconstituted GFP (green) and HRP (magenta) in third instar larvae of indicated genotypes (quantified in F). Neuronal Tnc influences the rGFP signals in the synaptic teminals but not in motor neuron axons. Scale bars: 10 μm; 5 μm in details. The number of examined NMJs is indicated below each bar. Bars indicated mean ±SEM. \*\*\**p*<0.001. Genotypes: *α-Spec*^*spGFP11*^; *N*>*GFP1-10;α-Spec*^*spGFP11*^ *(BG380-Gal4/+;UAS-GFP1-10/+; α-Spec*^*spGFP11*^*); N*>*GFP1-10; tnc*^*ko*^ *α-Spec*^*spGFP11*^, *(BG380-Gal4/+;UAS-GFP1-10/+; α-Spec*^*spGFP11*^, *tnc*^*ko*^); *N*>*GFP1-10, tnc*^*RNAi*^; *α-Spec*^*spGFP11*^ *(BG380-Gal4/+;UAS-GFP1-10/UAS-tnc*^*RNAi*^; *α-Spec*^*spGFP11*^).

## REFERENCES

Astorga, C., Jorquera, R.A., Ramirez, M., Kohler, A., Lopez, E., Delgado, R., Cordova, A., Olguin, P., and Sierralta, J. (2016). Presynaptic DLG regulates synaptic function through the localization of voltage-activated Ca(2+) Channels. Scientific reports 6, 32132.

Bennett, V., and Lorenzo, D.N. (2013). Spectrin- and ankyrin-based membrane domains and the evolution of vertebrates. Curr Top Membr 72, 1–37.

Beumer, K., Matthies, H.J., Bradshaw, A., and Broadie, K. (2002). Integrins regulate DLG/FAS2 via a CaM kinase II-dependent pathway to mediate synapse elaboration and stabilization during postembryonic development. Development 129, 3381–3391.

Brand, A.H., and Perrimon, N. (1993). Targeted gene expression as a means of altering cell fates and generating dominant phenotypes. Development 118, 401–415.

Budnik, V., Gorczyca, M., and Prokop, A. (2006). Selected methods for the anatomical study of Drosophila embryonic and larval neuromuscular junctions. Int Rev Neurobiol 75, 323–365.

Carlson, S.S., Valdez, G., and Sanes, J.R. (2010). Presynaptic calcium channels and alpha3-integrins are complexed with synaptic cleft laminins, cytoskeletal elements and active zone components. Journal of neurochemistry 115, 654–666.

Chavis, P., and Westbrook, G. (2001). Integrins mediate functional pre- and postsynaptic maturation at a hippocampal synapse. Nature 411, 317–321.

D’Este, E., Kamin, D., Gottfert, F., El-Hady, A., and Hell, S.W. (2015). STED nanoscopy reveals the ubiquity of subcortical cytoskeleton periodicity in living neurons. Cell reports 10, 1246–1251.

Featherstone, D.E., Davis, W.S., Dubreuil, R.R., and Broadie, K. (2001). Drosophila alpha- and beta-spectrin mutations disrupt presynaptic neurotransmitter release. J Neurosci 21, 4215–4224.

Feinberg, E.H., Vanhoven, M.K., Bendesky, A., Wang, G., Fetter, R.D., Shen, K., and Bargmann, C.I. (2008). GFP Reconstitution Across Synaptic Partners (GRASP) defines cell contacts and synapses in living nervous systems. Neuron 57, 353–363.

Fouquet, W., Owald, D., Wichmann, C., Mertel, S., Depner, H., Dyba, M., Hallermann, S., Kittel, R.J., Eimer, S., and Sigrist, S.J. (2009). Maturation of active zone assembly by Drosophila Bruchpilot. J Cell Biol 186, 129–145.

Fraichard, S., Bouge, A.L., Chauvel, I., and Bouhin, H. (2006). Tenectin, a novel extracellular matrix protein expressed during Drosophila melanogaster embryonic development. Gene Expr Patterns 6, 772–776.

Garcia-Caballero, A., Zhang, F.X., Hodgkinson, V., Huang, J., Chen, L., Souza, I.A., Cain, S., Kass, J., Alles, S., Snutch, T.P., et al. (2018). T-type calcium channels functionally interact with spectrin (alpha/beta) and ankyrin B. Mol Brain 11, 24.

Goodman, S.R. (1999). Discovery of nonerythroid spectrin to the demonstration of its key role in synaptic transmission. Brain Res Bull 50, 345–346.

Gratz, S.J., Goel, P., Bruckner, J.J., Hernandez, R.X., Khateeb, K., Macleod, G.T., Dickman, D., and O’Connor-Giles, K.M. (2019). Endogenous tagging reveals differential regulation of Ca(2+) channels at single AZs during presynaptic homeostatic potentiation and depression. J Neurosci.

Gratz, S.J., Rubinstein, C.D., Harrison, M.M., Wildonger, J., and O’Connor-Giles, K.M. (2015). CRISPR-Cas9 Genome Editing in Drosophila. Curr Protoc Mol Biol 111, 31 32 31–20.

Guzman, G.A., Guzman, R.E., Jordan, N., and Hidalgo, P. (2019). A Tripartite Interaction Among the Calcium Channel alpha1- and beta-Subunits and F-Actin Increases the Readily Releasable Pool of Vesicles and Its Recovery After Depletion. Front Cell Neurosci 13, 125.

Han, B., Zhou, R., Xia, C., and Zhuang, X. (2017). Structural organization of the actin-spectrin-based membrane skeleton in dendrites and soma of neurons. Proc Natl Acad Sci U S A 114, E6678–E6685.

He, J., Zhou, R., Wu, Z., Carrasco, M.A., Kurshan, P.T., Farley, J.E., Simon, D.J., Wang, G., Han, B., Hao, J., et al. (2016). Prevalent presence of periodic actin-spectrin-based membrane skeleton in a broad range of neuronal cell types and animal species. Proc Natl Acad Sci U S A 113, 6029–6034.

Huang, Z., Shimazu, K., Woo, N.H., Zang, K., Muller, U., Lu, B., and Reichardt, L.F. (2006). Distinct roles of the beta 1-class integrins at the developing and the mature hippocampal excitatory synapse. J Neurosci 26, 11208–11219.

Jan, L.Y., and Jan, Y.N. (1976). Properties of the larval neuromuscular junction in Drosophila melanogaster. J Physiol 262, 189–214.

Khanna, M.R., Mattie, F.J., Browder, K.C., Radyk, M.D., Crilly, S.E., Bakerink, K.J., Harper, S.L., Speicher, D.W., and Thomas, G.H. (2015). Spectrin tetramer formation is not required for viable development in Drosophila. J Biol Chem 290, 706–715.

Khanna, R., Li, Q., Bewersdorf, J., and Stanley, E.F. (2007a). The presynaptic CaV2.2 channel-transmitter release site core complex. The European journal of neuroscience 26, 547–559.

Khanna, R., Zougman, A., and Stanley, E.F. (2007b). A proteomic screen for presynaptic terminal N-type calcium channel (CaV2.2) binding partners. J Biochem Mol Biol 40, 302–314.

Kittel, R.J., Wichmann, C., Rasse, T.M., Fouquet, W., Schmidt, M., Schmid, A., Wagh, D.A., Pawlu, C., Kellner, R.R., Willig, K.I., et al. (2006). Bruchpilot promotes active zone assembly, Ca2+ channel clustering, and vesicle release. Science 312, 1051–1054.

Machnicka, B., Czogalla, A., Hryniewicz-Jankowska, A., Boguslawska, D.M., Grochowalska, R., Heger, E., and Sikorski, A.F. (2014). Spectrins: a structural platform for stabilization and activation of membrane channels, receptors and transporters. Biochimica et biophysica acta 1838, 620–634.

Macleod, G.T. (2012). Forward-filling of dextran-conjugated indicators for calcium imaging at the Drosophila larval neuromuscular junction. Cold Spring Harb Protoc 2012, 791–796.

Macleod, G.T., Chen, L., Karunanithi, S., Peloquin, J.B., Atwood, H.L., McRory, J.E., Zamponi, G.W., and Charlton, M.P. (2006). The Drosophila cacts2 mutation reduces presynaptic Ca2+ entry and defines an important element in Cav2.1 channel inactivation. The European journal of neuroscience 23, 3230–3244.

Macleod, G.T., Marin, L., Charlton, M.P., and Atwood, H.L. (2004). Synaptic vesicles: test for a role in presynaptic calcium regulation. J Neurosci 24, 2496–2505.

McMahan, U.J., Horton, S.E., Werle, M.J., Honig, L.S., Kroger, S., Ruegg, M.A., and Escher, G. (1992). Agrin isoforms and their role in synaptogenesis. Curr Opin Cell Biol 4, 869–874.

Mosca, T.J., Hong, W., Dani, V.S., Favaloro, V., and Luo, L. (2012). Trans-synaptic Teneurin signalling in neuromuscular synapse organization and target choice. Nature 484, 237–241.

Nishimune, H., Sanes, J.R., and Carlson, S.S. (2004). A synaptic laminin-calcium channel interaction organizes active zones in motor nerve terminals. Nature 432, 580–587.

Orlacchio, A., Calabresi, P., Rum, A., Tarzia, A., Salvati, A.M., Kawarai, T., Stefani, A., Pisani, A., Bernardi, G., Cianciulli, P., et al. (2007). Neuroacanthocytosis associated with a defect of the 4.1R membrane protein. BMC Neurol 7, 4.

Park, Y.K., and Goda, Y. (2016). Integrins in synapse regulation. Nat Rev Neurosci 17, 745–756.

Peng, I.F., and Wu, C.F. (2007). Drosophila cacophony channels: a major mediator of neuronal Ca2+ currents and a trigger for K+ channel homeostatic regulation. J Neurosci 27, 1072–1081.

Pfeiffer, B.D., Ngo, T.T., Hibbard, K.L., Murphy, C., Jenett, A., Truman, J.W., and Rubin, G.M. (2010). Refinement of tools for targeted gene expression in Drosophila. Genetics 186, 735–755.

Phillips, G.R., Huang, J.K., Wang, Y., Tanaka, H., Shapiro, L., Zhang, W., Shan, W.S., Arndt, K., Frank, M., Gordon, R.E., et al. (2001). The presynaptic particle web: ultrastructure, composition, dissolution, and reconstitution. Neuron 32, 63–77.

Pielage, J., Fetter, R.D., and Davis, G.W. (2005). Presynaptic spectrin is essential for synapse stabilization. Curr Biol 15, 918–928.

Qu, Y., Hahn, I., Webb, S.E., Pearce, S.P., and Prokop, A. (2017). Periodic actin structures in neuronal axons are required to maintain microtubules. Molecular biology of the cell 28, 296–308.

Ren, X., Sun, J., Housden, B.E., Hu, Y., Roesel, C., Lin, S., Liu, L.P., Yang, Z., Mao, D., Sun, L., et al. (2013). Optimized gene editing technology for Drosophila melanogaster using germ line-specific Cas9. Proc Natl Acad Sci U S A 110, 19012–19017.

Ribeiro, S.A., D’Ambrosio, M.V., and Vale, R.D. (2014). Induction of focal adhesions and motility in Drosophila S2 cells. Molecular biology of the cell 25, 3861–3869.

Rohrbough, J., Grotewiel, M.S., Davis, R.L., and Broadie, K. (2000). Integrin-mediated regulation of synaptic morphology, transmission, and plasticity. J Neurosci 20, 6868–6878.

Ruiz-Canada, C., and Budnik, V. (2006). Synaptic cytoskeleton at the neuromuscular junction. Int Rev Neurobiol 75, 217–236.

Shurer, C.R., Kuo, J.C., Roberts, L.M., Gandhi, J.G., Colville, M.J., Enoki, T.A., Pan, H., Su, J., Noble, J.M., Hollander, M.J., et al. (2019). Physical Principles of Membrane Shape Regulation by the Glycocalyx. Cell 177, 1757–1770 e1721.

Sidenstein, S.C., D’Este, E., Bohm, M.J., Danzl, J.G., Belov, V.N., and Hell, S.W. (2016). Multicolour Multilevel STED nanoscopy of Actin/Spectrin Organization at Synapses. Scientific reports 6, 26725.

Smith, L.A., Wang, X., Peixoto, A.A., Neumann, E.K., Hall, L.M., and Hall, J.C. (1996). A Drosophila calcium channel alpha1 subunit gene maps to a genetic locus associated with behavioral and visual defects. J Neurosci 16, 7868–7879.

Stewart, B.A., Atwood, H.L., Renger, J.J., Wang, J., and Wu, C.F. (1994). Improved stability of Drosophila larval neuromuscular preparations in haemolymph-like physiological solutions. J Comp Physiol A 175, 179–191.

Sulkowski, M.J., Han, T.H., Ott, C., Wang, Q., Verheyen, E.M., Lippincott-Schwartz, J., and Serpe, M. (2016). A Novel, Noncanonical BMP Pathway Modulates Synapse Maturation at the Drosophila Neuromuscular Junction. PLoS genetics 12, e1005810.

Sunderland, W.J., Son, Y.J., Miner, J.H., Sanes, J.R., and Carlson, S.S. (2000). The presynaptic calcium channel is part of a transmembrane complex linking a synaptic laminin (alpha4beta2gamma1) with non-erythroid spectrin. J Neurosci 20, 1009–1019.

Syed, Z.A., Bouge, A.L., Byri, S., Chavoshi, T.M., Tang, E., Bouhin, H., van Dijk-Hard, I.F., and Uv, A. (2012). A luminal glycoprotein drives dose-dependent diameter expansion of the Drosophila melanogaster hindgut tube. PLoS genetics 8, e1002850.

Venken, K.J., He, Y., Hoskins, R.A., and Bellen, H.J. (2006). P[acman]: a BAC transgenic platform for targeted insertion of large DNA fragments in D. melanogaster. Science 314, 1747–1751.

Wagh, D.A., Rasse, T.M., Asan, E., Hofbauer, A., Schwenkert, I., Durrbeck, H., Buchner, S., Dabauvalle, M.C., Schmidt, M., Qin, G., et al. (2006). Bruchpilot, a protein with homology to ELKS/CAST, is required for structural integrity and function of synaptic active zones in Drosophila. Neuron 49, 833–844.

Wang, Q., Han, T.H., Nguyen, P., Jarnik, M., and Serpe, M. (2018). Tenectin recruits integrin to stabilize bouton architecture and regulate vesicle release at the Drosophila neuromuscular junction. Elife 7.

Xu, K., Zhong, G., and Zhuang, X. (2013). Actin, spectrin, and associated proteins form a periodic cytoskeletal structure in axons. Science 339, 452–456.

Zhong, G., He, J., Zhou, R., Lorenzo, D., Babcock, H.P., Bennett, V., and Zhuang, X. (2014). Developmental mechanism of the periodic membrane skeleton in axons. Elife 3.

